# Vitamins D3 and D2 have marked but different global effects on gene expression in a rat oligodendrocyte precursor cell line

**DOI:** 10.1101/730168

**Authors:** Manuela Mengozzi, Andrew Hesketh, Giselda Bucca, Pietro Ghezzi, Colin P. Smith

## Abstract

Vitamin D deficiency increases the risk of developing multiple sclerosis (MS) but there is uncertainty about what dose and form of vitamin D could improve the clinical course of MS. The mechanisms underlying the effects of vitamin D in MS are not clear. Vitamin D3 increases the rate of differentiation of primary oligodendrocyte precursor cells (OPCs), suggesting that it might help remyelination in addition to modulating the immune response. Here we analyzed the transcriptome of differentiating rat CG4 OPCs treated with vitamin D2 or with D3 at 24 h and 72 h following onset of differentiation. Differentiation alone changed the expression of about 10% of the genes at 72 h compared to 24 h. Vitamin D2 and D3 exerted different effects on gene expression, with D3 influencing 1,272 genes and D2 574 at 24 h. The expression of the vast majority of these genes was either not changed in differentiating cells not exposed to vitamin D or followed the same trajectory as the latter. D3-repressed genes were enriched for gene ontology categories including transcription factors and the Notch pathway, while D3-induced genes were enriched for the Ras pathway. These findings should help to identify mechanisms mediating D3 action in MS.

## Introduction

There is evidence that vitamin D deficiency is associated with increased risk of developing multiple sclerosis (MS) and with an increased rate of disease progression^1,2^. However, the therapeutic efficacy of vitamin D supplementation in MS patients is still an open question, and interventional trials have reported inconclusive results^3^.

Vitamin D is produced in the skin by the action of ultraviolet light. After two successive hydroxylations, the 1,25(OH)D active form binds to the intracellular vitamin D receptor (VDR) and, in complex with the retinoid X receptor (RXR), regulates hundreds of genes through the vitamin D response element (VDRE)^1,4^.

Vitamin D can also be assimilated from the diet or as a supplement. Both the plant/fungus-derived vitamin D2, ergocalciferol, and the animal-derived D3 form, cholecalciferol, are available as nutritional supplements; both can be hydroxylated into their active forms and bind VDR with similar affinity^5^, but there are differences in their catabolism and in their binding affinity to vitamin D binding protein (DBP), the major vitamin D transport protein in blood, with vitamin D2 binding DBP with lower affinity and being catabolised more quickly^1,4^.

Studies comparing vitamin D2 and D3 for their ability to raise and maintain circulating levels of 25(OH)D, the serum marker of systemic vitamin D levels, have shown that vitamin D2 is less effective than D3 when given as single bolus^6^; however, results following daily administration of D2 or D3 are controversial, with clinical trials showing higher efficacy of D3^7,8^ or equal efficacy^9^. In a recent meta-analysis, vitamin D2 and D3 were found equally effective in raising vitamin D levels in infants^10^. It has been suggested that vitamin D2 supplementation reduces circulating vitamin D3 levels; however, a recent randomized controlled trial showed that the reciprocal is also true^11^. The general consensus is that vitamin D3 is the preferred form to be administered^3^, but few studies have compared D2 and D3 in terms of their direct cellular effects under controlled conditions.

The mechanisms mediating the protective action of vitamin D in MS are not fully characterized. In addition to regulating calcium homeostasis, vitamin D is immunomodulatory and anti-inflammatory^1,2,12^. In the last twenty years, many studies have reported effects on the brain; vitamin D inhibits neuro-inflammation, is neuroprotective and neurotrophic^1,13–15^. VDR is present in neurons and glial cells^16,17^. Vitamin D acts directly on brain cells by promoting the differentiation of neural stem cells (NSCs) into neurons and oligodendrocytes (OLs), and inducing the expression of growth factors, such as nerve growth factor (NGF), in neurons and OLs^16,18^.

The regenerative and neurotrophic effects of vitamin D, together with the presence of VDR in OLs, suggest that it might have a myelinating action, which might contribute to its protective effects in MS. In this respect, de la Fuente *et al*. found that vitamin D increased the differentiation and maturation of primary oligodendrocyte progenitor cells (OPCs)^19^.

Rat central glia-4 (CG4) OPCs are considered a good model of myelination *in vitro*; the cells are maintained at the precursor stage by culture with growth factors (GFs), and can be differentiated into OLs by withdrawal of GFs and culture in differentiation medium (DM)^20–22^. Interestingly, Baas *et al*. found that CG4 cells express VDR, as primary OPCs, and respond to vitamin D by increasing expression of VDR and of NGF; however, an effect on myelin proteins was not found, leading to the conclusion that vitamin D does not positively affect the differentiation of these cells^16^.

To better understand the molecular mechanisms by which vitamin D acts on OLs, we analyzed the gene expression profile of rat CG4 OPCs in response to vitamin D, and compared vitamin D2 with D3.

## Results

### Set up and experimental treatment

We first established the appropriate experimental conditions for studying the effect of vitamin D on the gene expression profile of differentiating CG4 cells. In a previous study, Baas *et al*. found that VDR is present in CG4 OL precursors and its levels are increased 2-fold after culture for two days in differentiating conditions^16^; we therefore hypothesized that a pre-incubation in differentiation medium (DM) for two days before treatment with vitamin D might increase the basal levels of VDR and the response to vitamin D. Since VDR is induced by vitamin D3^16^, we monitored basal and vitamin D3-induced VDR levels in different culture conditions. The dose of 100 nM vitamin D was chosen based on previous studies^16,19^.

Cells were plated, left to adhere overnight, then medium was changed to DM and cells were treated with vitamin D3 added at the same time as DM (Fig. 1a) or after pre-incubation in DM for 48 h (Fig. 1b). VDR mRNA was measured after 24 h or 72 h of incubation with vitamin D3 in both conditions. A 48-h pre-incubation in DM increased basal levels of VDR and vitamin D3-induced VDR levels at 24 h and 72 h (Fig. 1). Of note, VDR expression was induced by vitamin D3 at 24 h only after cells were pre-differentiated for 48 h.

**Figure 1.**
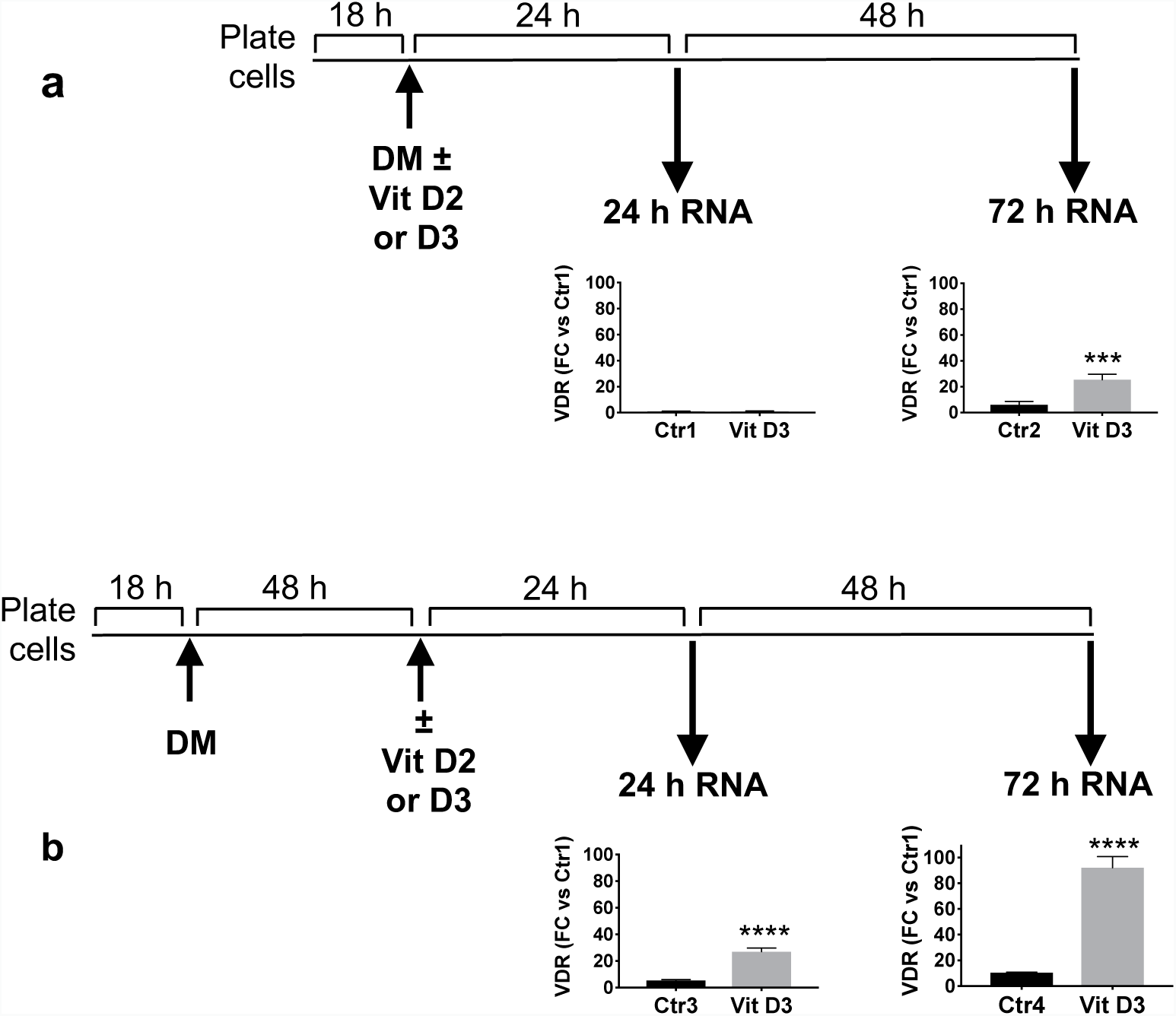
Experimental design of cell differentiation and treatment with vitamin D3 (Vit D3). Cells were plated and cultured overnight before switching to differentiation medium (DM). Vit D3 (100 nM) or vehicle (Ctr) was added simultaneously with DM (**a**) or after 48 h (**b**). VDR gene expression was measured by qPCR 24 h or 72 h after vitamin D3 or vehicle addition. Data are expressed as fold change (FC) versus ctr1 and are the mean ± SD of four replicates. *** P < 0.001, **** P < 0.0001 by Student’s t-test. QPCR analysis was not conducted on D2-treated cultures.

Therefore, the experimental design in Fig. 1b was used for the transcriptome analysis. After 48 h of pre-differentiation in DM, cells were treated with either vitamin D2 or D3; RNA was isolated following incubation for a further 24 h and 72 h and analyzed using whole genome DNA microarrays. The DNA microarray design uses unique oligonucleotide probes to quantify the abundance of gene transcripts containing the complementary sequences, and can include more than one probe per gene. In the presentation of the results below we make the distinction between probes and genes when necessary, and occasionally use the term “probes-genes” when no distinction is required.

### Effect of vitamin D2 and D3 on gene expression profile

Overall, addition of either vitamin D2 or D3 to differentiating cells significantly affected gene expression, an effect that was more marked at 72 h (Fig. 2). Out of 15,247 probes, corresponding to 12,261 genes (ArrayExpress Accession number E-MTAB-8098) about 30% were affected by either vitamin D2 or D3 at 72 h, and 10% at 24 h (Fig. 2a).

**Figure 2.**
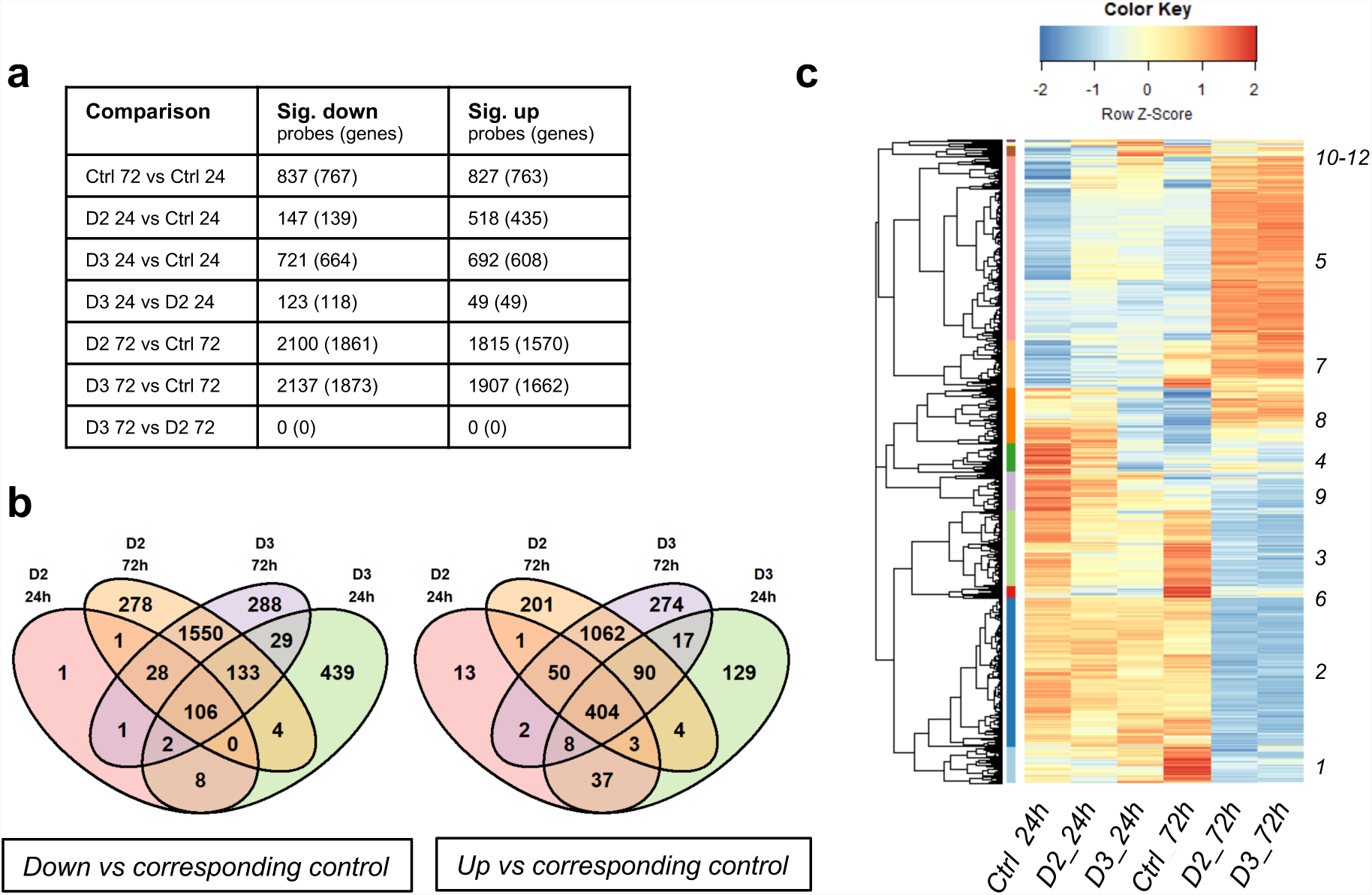
Probes-genes regulated by vitamin D2, vitamin D3 or differentiation alone at 24 and 72 h. (**a**) Values represent numbers of probes detecting differentially expressed genes compared with their respective control (adj.P.Val ≤ 0.05, testing significance relative to a 10%-fold change in expression using LIMMA TREAT^52^. The Venn diagrams (**b**) show the number of probes specifically down-regulated (left) or up-regulated (right) by each vitamin or both. The Ctrl 72 h vs Ctrl 24 h data is not represented in the Venn diagrams. (**c**) Hierarchical cluster analysis and heatmap of all the probes significantly different in any of the comparisons listed in panel **a** (5,565 probes). All the probes-genes are listed in Supplementary File S1, where information about the respective clusters is included.

The numbers of probes-genes affected by vitamin D2 and D3 were broadly similar at 72 h, with most differentially expressed genes regulated by both D2 and D3 (Fig. 2a, b). However, the shorter-term response at 24 h showed a more pronounced response to vitamin D3, with notably larger numbers of probes-genes down-regulated compared with vitamin D2 (721 versus 147 probes, corresponding to 664 versus 139 genes respectively; Fig. 2a). At 24 h, most genes regulated by D2 were also regulated by D3, with some other genes regulated by D3 only (Fig. 2b).

To visualize the global effect of vitamin D2 and D3 in relation to untreated cells, we performed a hierarchical cluster analysis of the 5,565 probes differentially regulated when comparing vitamin D2 or D3 with their respective controls at both time points, as well as untreated controls at 72 h versus 24 h. The results are shown as a heatmap in Fig. 2c.

By comparing untreated control cells (Ctrl) at 72 h with Ctrl at 24 h, we can see that differentiation alone noticeably affects gene expression. The actual number of probes-genes regulated in Ctrl 72 h versus Ctrl 24 h is reported in Fig. 2a. Figure 2c also shows that a number of genes with low expression in control cells are up-regulated by both vitamin D2 and D3 at 24 h, an effect even more marked than that of differentiation at 72 h (e.g. cluster 5 in Fig. 2c). Likewise, some of the genes that have a high expression in 24 h Ctrl cells seem to be down-regulated by vitamin D2 and D3 at 24 h, an effect similar to differentiation alone (e.g. cluster 9 in Fig. 2c). This trend is confirmed at 72 h, where vitamin D2 and D3 have a more marked effect than at 24 h.

### Effect of differentiation

To analyze the effect of vitamin D in relation to the effect of differentiation, we focused on the genes changed by culture in DM alone. As reported in Fig. 2a, 837 probes (767 genes) were down-regulated and 827 (763 genes) were up-regulated in untreated cells (Ctrl) at 72 h compared to Ctrl at 24h. The functional enrichment analysis of the 767 down-regulated and of the 763 up-regulated genes in the 72 h versus 24 h control cells is reported in Fig. 3a. This reveals that most of the down-regulated genes were associated with the cell cycle and the up-regulated genes were significantly (p.adjust ≤ 0.05) enriched for Gene Ontology Biological Process (GO-BP) categories including “cell maturation” (p.adjust 7.77E-03), “axon ensheathment” (p.adjust 2.28E-03), “myelination” (p.adjust 6.75E-03); this is consistent with the fact that CG4 cells in DM differentiate over time^22,23^.

**Figure 3.**
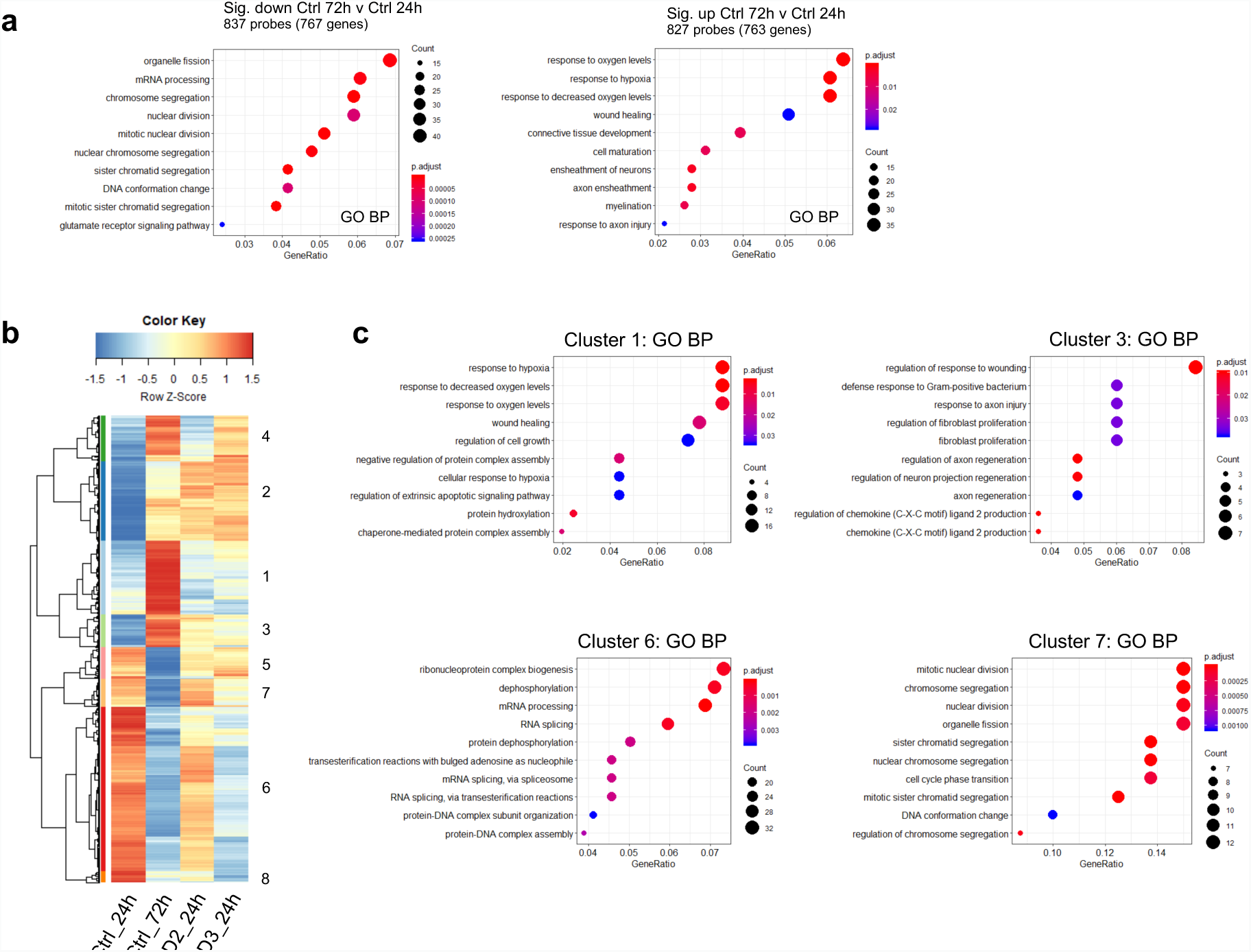
Functional enrichment analysis of the genes regulated during differentiation alone and effect of vitamin D2 and D3 on these genes. (**a**) The enriched Gene Ontology Biological Processes (GO-BPs) among the 767 down-regulated (left) and 763 up-regulated (right) genes, shown in Fig. 2A as “Ctrl 72 vs Ctrl 24”, are listed. The size of the dots is proportional to the number of genes and the color represents the corrected p-value (p.adjust), as indicated in the legend. (**b**) Cluster analysis and heat map of the expression data for the probes corresponding to the genes in (**a**) (837 down-regulated and 827 up-regulated by differentiation alone, Fig. 2a); the expression signal at 24 h and 72 h in untreated cells and at 24 h in D2- and D3-treated cells is shown, to visualize the effect of D2 and D3 versus their control at 24 h and the effect of differentiation alone at 72 h versus 24 h. (**c**) Enriched GO-BP terms corresponding to some of the clusters. All the probes-genes included in the clusters and in the GO-BPs are listed in Supplementary File S2.

To visualize the effect of vitamin D2 and D3 at 24 h on the probes changed by differentiation alone (837 and 827 respectively, as mentioned above), we performed a hierarchical cluster analysis (Fig. 3c) of the expression abundance of these probes in untreated cells at 24 and 72 h (highlighting the effect of differentiation alone) and in vitamin D2- and D3-treated cells at 24 h (highlighting the effect of each vitamin, that can be visually compared to the effect of differentiation alone).

The clusters 1-4 in Fig. 3b comprise all the probes corresponding to the genes increased by differentiation alone. Both vitamins, and in particular vitamin D3, preferentially increase the expression of these genes at 24 h.

The same trend was confirmed for the down-regulated genes (clusters 5-8 in Fig. 3b), where the preferential downregulating effect of vitamin D3 compared to D2 was more marked. Functional enrichment analysis of the down-regulated clusters 6 and 7 in Fig. 3c identified GO-BP categories associated with cell cycle, whereas “axon regeneration” (p.adjust 3.73E-02) was one of the GO-BP terms significantly enriched among the clusters increased by differentiation and by vitamin D2 and D3 (cluster 3, Fig. 3c).

### Analysis of vitamin D-regulated genes in relation to differentiation

A more specific analysis of the effects of vitamin D on gene expression in relation to the change in the gene expression profile induced by differentiation alone (Ctrl 72 h versus Ctrl 24 h) is shown in Fig. 4.

**Figure 4.**
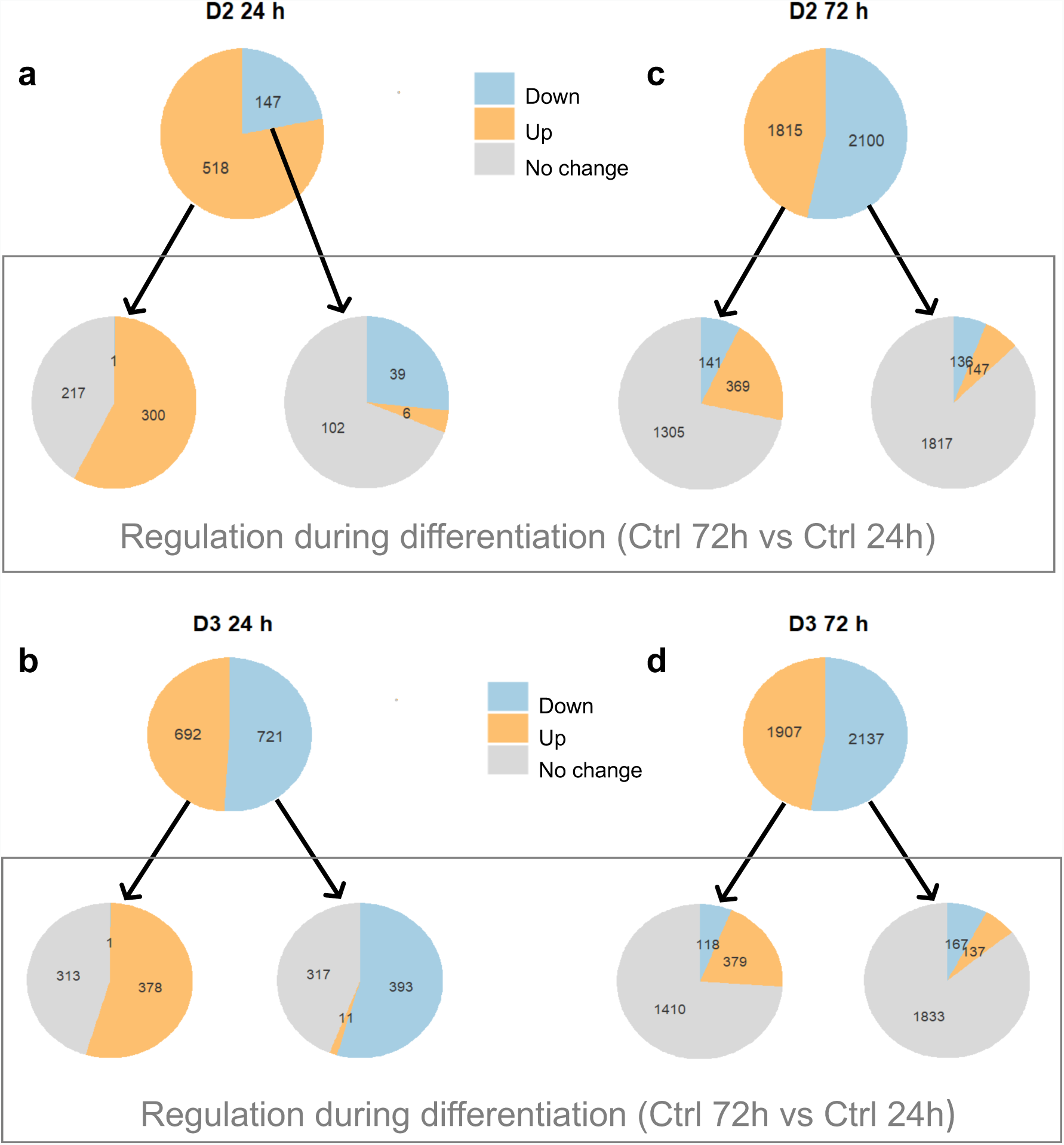
Genes regulated by D2 and D3 at 24 h and 72 h and relative effect of differentiation alone. The numbers of significant probes are indicated in each pie chart. The top pie charts in all panels indicate the number of probes that are up- or down-regulated by vitamin D2 (**a** and **c**) or D3 (**b** and **d**) compared to the control at the same time point (**a** and **b**, 24 h; **c** and **d**, 72 h). The bottom pie charts of all panels describe how these genes are regulated by differentiation (defined by comparing Ctrl 72h vs Ctrl 24h). Differentially regulated genes were obtained according to the same criteria described in the legend to Fig. 2A. All the probes-genes are included in Supplementary File S1, where the membership of the pie chart groups is reported.

More than half of the genes up-regulated (in orange in the figure) at 24 h by vitamin D2 (a) or D3 (b) were also induced by differentiation, with the remaining genes (in grey in the figure) not affected by differentiation alone. Only one gene (spondin 1, Supplementary File S1) induced by vitamin D2 or D3 at 24 h was changed in the opposite direction by differentiation.

Of the genes that were down-regulated at 24 h (in blue in Fig. 4) by vitamin D2 (a) or D3 (b), most genes were down-regulated (in blue in the bottom pie charts of the two panels) or unaffected (in grey in the bottom pie charts of the panels) by differentiation. Only few vitamin D-down-regulated probes (six for D2 and eleven for D3) were changed in the opposite direction by differentiation. In summary, a 24 h exposure to vitamin D has either a unique effect for specific genes or an effect on gene expression similar to that of differentiation.

A similar analysis was performed at 72 h (Fig. 4c and d). At this time point, the overlap between vitamin D- and differentiation-regulated genes is smaller, and most of the effect on the gene expression profile is unique to vitamin D.

### Genes uniquely regulated by vitamin D3 and not by D2

Because vitamin D3 had a more marked effect on gene expression at 24 h than D2 (Fig. 2a, Fig. 4a and b), we analyzed the genes uniquely regulated by D3 (but not by D2) at 24 h and their behavior in relation to differentiation. As shown in Fig. 5, all the genes uniquely regulated by vitamin D3 were either not changed by differentiation in absence of vitamin D, or followed the same trajectory as differentiation, with only five probes down-regulated by D3 but up-regulated by differentiation.

**Figure 5.**
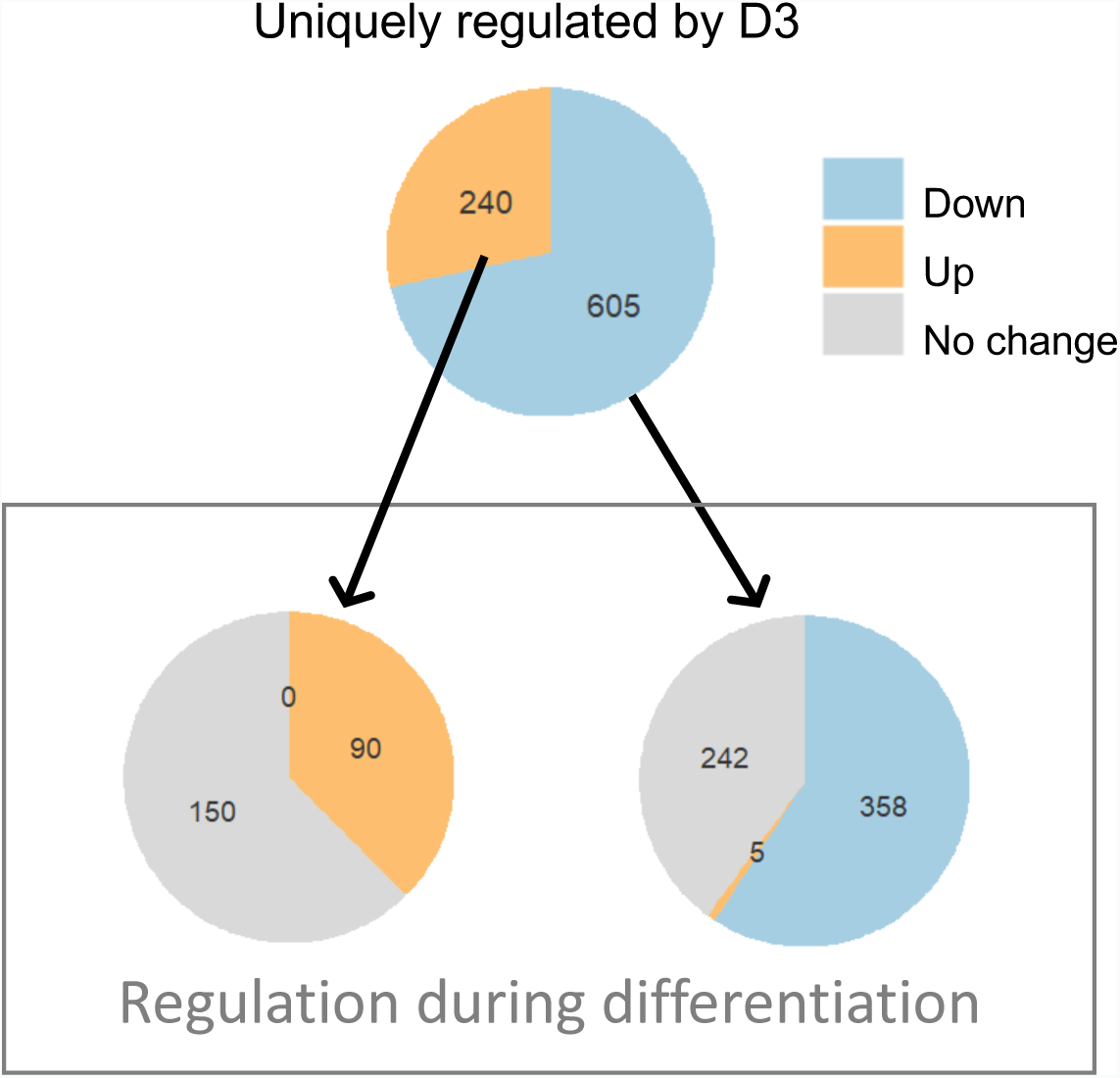
Genes uniquely regulated by vitamin D3 at 24 h and relative effect of differentiation alone. Numbers in the top pie charts refer to probes up-regulated (orange) or down-regulated (blue) by D3 versus control but not by D2 versus control. The list of the probes-genes is reported in Supplementary File S1.

### Gene expression profile in oligodendrocyte cells is more greatly affected by vitamin D3 than D2

To analyze the differential effect on gene expression of vitamin D3 compared to D2, we focused on the probes-genes uniquely regulated by vitamin D3 (and not by vitamin D2) at 24 h. At this time point, vitamin D3 uniquely down-regulated 605 probes and up-regulated 240 probes, as shown in Fig. 5a.

Functional enrichment analysis of the genes corresponding to the probes significantly decreasing in abundance after 24 h in response to either vitamin D2 (n=147, Fig. 2a and 4a) or D3 (n=721, Fig. 2a and 4b) and of those uniquely down-regulated by D3 (n=605; Fig. 5) indicated that the two forms of vitamin D produce notably different early effects on oligodendrocyte cells (Supplementary Fig. S1, Supplementary File S3). Interestingly, the 605 genes uniquely down-regulated by D3 at 24 h were significantly enriched for GO terms associated with transcription factor complex activity (“transcription factor complex”, p.adjust 3.59E-03; “transcription factor activity”, p.adjust 8.29E-04) suggesting that the two forms of vitamin D might exert different effects on transcriptional regulation by these complexes. Other functions similarly and potentially more adversely affected by D3 compared to D2 at 24 h include “RNA splicing” (p.adjust 3.07E-02), “axonogenesis” (p.adjust 7.73E-03), Toll-like receptor (TLR) cascade pathways (e.g. “TLR3 cascade”, p.adjust 4.01E-03), “MAPK signaling pathways” (p.adjust 2.25E-02) and “Notch signaling pathway” (p.adjust 2.71E-02).

Functional enrichment analysis of the genes corresponding to the probes significantly increased at 24 h by vitamin D2 (n=518, Fig. 2a and 4a) or D3 (n=692; Fig. 2a and 4b) and of those uniquely up-regulated by D3 (n=240, Fig. 5) is reported in Supplementary Fig. S2 and Supplementary File S3. The genes uniquely up-regulated by D3 were enriched for the GO term “GTP binding” (p.adjust 4.11E-02) whereas analysis of all the genes up-regulated by vitamin D3 showed an enrichment of the Ras signal transduction pathway (p.adjust 3.46E-02).

## Discussion

The main finding of this study is that vitamin D2 and D3 exert markedly different effects on gene expression in differentiating OPCs. Although there is some overlap in the genes they regulate, D3 affects a larger repertoire of genes. In particular, we identified some genes that are specifically influenced by D2 only (31 at 24 h and 66 at 72 h) or D3 only (605 at 24 h and 240 at 72 h), showing that, in this experimental model, D2 and D3 (or their respective derivatives) might behave as distinct molecular entities with different biological activities.

*In vivo*, studies where D2 and D3 are compared are likely to mainly reflect differences in their pharmacokinetics. This should not be an issue in an *in vitro* model that should reflect the direct effect of the molecule tested, and our results are likely to relate to differences in the biological activity of the two forms of vitamin D. Although, reportedly, vitamin D2 and D3 have similar affinities for VDR^4,5^, they may have different affinity for other vitamin D binding proteins. In fact, the action of vitamin D is mediated by other receptors and binding proteins such as the retinoid-related orphan receptors (RORs) and Erp57, a membrane steroid-binding protein related to protein disulfide isomerase^24–26^. These additional receptors may have important biological roles in the central nervous system, and the protective effect of D3 on axonal growth has been linked to activation of membrane-associated rapid response steroid-binding receptor (MARRS)/Erp57^27^.

Vitamin D3 and D2 both up-regulated VDR expression, which was also up-regulated during OPC differentiation, in agreement with a previous report^16^. Both vitamins regulated the expression of many genes that were either not changed in differentiating cells not exposed to vitamin D or followed the same trajectory as the latter; only very few genes followed the opposite expression trajectory, suggesting that in this system, as in primary OLs^19^, vitamin D promotes rather than antagonizing differentiation.

Although at 72 h both vitamin D2 and D3 had similar effects on gene expression, at 24 h vitamin D3 was more potent, changing the expression of a substantially higher number of genes. The preferential action of D3 versus D2 was particularly evident when considering the down-regulated genes; only about one fourth of the genes down-regulated by D3 were also down-regulated by D2 at 24 h, whereas most of the genes down-regulated by D2 were also down-regulated by D3, as shown in Fig. 2. The higher efficacy of vitamin D3 compared to D2 in increasing myelination had been previously reported in a rat model of peripheral nerve injury^28^, but no studies had directly compared vitamin D2 and D3 *in vitro* on OL differentiation. It should be noted, however, that we did not observe any direct effect of vitamin D on the expression of myelin genes, in agreement with previous studies^16^, highlighting a limitation of this experimental model.

Functional enrichment analysis of the genes uniquely down-regulated by D3 (unchanged by D2) at 24 h highlighted an over-representation of transcription factors; among these, SRY-related HMG-Box Gene 4 (Sox4) and transcription factor EB (Tfeb), whose high expression has been previously associated with inhibition of myelination^29,30^. In addition, the KEGG MAPK signaling pathway was overrepresented; of note, we previously reported that in this model inhibition of ERK1/2 MAPK increases OL differentiation^22^. Fibroblast growth factor receptor 2 (Fgfr2) and platelet derived growth factor A (Pdgfa) were included in this pathway (Supplementary File S3); since both Fgf and Pdgf are needed to maintain CG4 OPCs in the undifferentiated state^20,22^, inhibition of Fgfr and Pdgf would promote differentiation.

Considering all the genes down-regulated by D3 at 24 h (and therefore possibly also by D2), enrichment of the Notch signaling pathway was identified. Activation of the Notch pathway inhibits OPC differentiation during development and is involved in the limited remyelination that characterizes MS^31,32^. Notch is a transmembrane receptor that is activated upon binding to several ligands, including members of the Jagged and Delta-like families; these ligands induce sequential cleavage of Notch, with an essential cleavage performed by a gamma-secretase, which contains presenilin^33^.

The ligands Jagged 1 (Jag1) and delta-like canonical Notch ligand 3 (Dll3) were inhibited by both vitamin D2 and D3; in addition, presenilin 1 (Psen1) was inhibited only by vitamin D3. The inhibitory effect of vitamin D3 on gamma-secretase activity had been previously reported in the context of Alzheimer’s disease^34^; in this respect, gamma-secretase, in addition to Notch, also cleaves the amyloid precursor protein (APP) to produce amyloid-beta. Inhibition of Notch ligands and of Psn1, and of other genes belonging to the Notch pathway (Supplementary File S3), suggests a differentiating effect of vitamin D on these cells. In addition, preferential inhibition of Psn1 by vitamin D3 and not D2 indicates again that D3 is more efficacious than D2. Interestingly, inhibition of Notch-signalling molecules, which play a role in cancer stem cell maintenance, is one of the mechanisms that mediate the anticancer properties of vitamin D^35,36^.

Functional enrichment analysis of all the genes up-regulated by D3 revealed overrepresentation of the GO-BP Ras protein signal transduction (Supplementary Fig. S2, Supplementary File S3). GTPases of the Ras family affect cell survival, proliferation and differentiation^37^, and among these Rras2 was recently found to be essential for OL differentiation^38^. Included in this pathway are also nerve growth factor (Ngf) and transforming growth factor beta 2 (Tgfb2), both reported to activate Ras-dependent signalling pathways and to promote OPC differentiation^39–43^. Vitamin D3-induced upregulation of Ngf in these cells was previously reported^16^. Of note, Rras2 and Ngf were also induced by D2, whereas Tgfb2 was uniquely induced by D3 (Supplementary File S3). Also included in this pathway were several Rab genes, such as Rab36, Rab9d, Rab6a, and Rab3d. Rab proteins, involved in vesicular transport, are up-regulated during OL differentiation, in particular when OL start synthesizing myelin^44^; interestingly Rab6a and Rab3d were uniquely up-regulated by D3 (GO Molecular Function (GO-MF) “GTP binding”, Supplementary Fig. S2; Supplementary File S3). In addition, vitamin D3-induced transcription factors (GO Cellular Component (GO-CC) “nuclear transcription factor complex”, Supplementary Fig. S2) included Sox2, whose role in OL differentiation was recently reported^45,46^ and CCAAT/enhancer binding protein beta (Cebpb), a VDR-responsive gene that controls the differentiation of myeloid leukaemia cells^47^.

Our identification of specific vitamin D3-regulated genes in OPCs could provide important insights to understand the mechanisms mediating the activity of vitamin D in demyelinating disease and point to mechanisms additional to those associated with immune regulation, the latter being a major research focus for the action of vitamin D in the context of MS^1^.

## Methods

### Cell culture

The rat CG4 OL precursor cell line (OPC), originally obtained from primary cultures of bipotential oligodendrocyte-type-2-astrocytes (O-2A)^20^, was cultured as previously reported ^22,23^. CG4 cells are maintained at the precursor stage by culture in growth medium (GM), consisting of Dulbecco’s modified Eagle medium (DMEM; Sigma-Aldrich) supplemented with biotin (10 ng/ml), bFGF (5 ng/ml), PDGF (1 ng/ml), N1 supplement (all from Sigma-Aldrich) and 30% B104-conditioned medium, obtained as previously reported^23^. The cells can be differentiated into mature OLs by withdrawal of growth factors (bFGF and PDGF) and of B104-conditioned medium and culture in differentiation medium (DM), consisting of DMEM-F12 (Invitrogen) supplemented with progesterone (3 ng/ml), putrescine (5 µg/ml), sodium selenite (4 ng/ml), insulin (12.5 µg/ml), transferrin (50 µg/ml), biotin (10 ng/ml), thyroxine (0.4 µg/ml) and glucose (3 g/l) (all from Sigma-Aldrich).

Cells were plated in poly-L-ornithine-coated 12-well plates (100,000 cells in 2 ml GM per well). After overnight culture, the cells were induced to differentiate by switching to DM. Cells were cultured for the indicated times; half of the medium was changed every other day. For vitamin D treatment, vitamin D2 (1,25-dihydroxyvitamin D2; Cayman Chemical) or vitamin D3 (1,25-dihydroxyvitamin D3; Sigma-Aldrich) were dissolved in ethanol (96%; Sigma-Aldrich) at 100 µM and then diluted in medium at the final concentration (100 nM). Vehicle (ethanol 0.1%) was added to untreated cells.

### RNA extraction

Each sample was lysed with 1 ml of QIAzol (QIAGEN). Total RNA was extracted using the miRNeasy system and protocol (QIAGEN). RNA concentration, purity and integrity were determined using a NanoDrop 1C (NanoDrop Technologies) spectrophotometer and an Agilent 4200 TapeStation (Agilent Technologies). All samples had a A260/A280 ratio > 1.8 and RNA Integrity Number > 8.2.

### RT-qPCR

Reverse transcription (RT) and real time quantitative PCR (qPCR) were carried out as reported^22^, using TaqMan® gene expression assays (Applied Biosystems/Thermo Fisher Scientific). Results were normalized to HPRT1 expression (reference gene) and expressed as fold change (FC) versus one of the control samples (as indicated), chosen as the calibrator.

### Microarrays

Biological quadruplicate samples were analyzed for each treatment group. In total, 24 arrays were analyzed: 4 untreated, 4 vitamin D2 and 4 vitamin D3 at each time point (following, respectively, 24 h and 72 h of treatment). Total RNA (200 ng) was amplified by *in vitro* transcription using the Low Input Quick Amp One color kit, incorporating Cy3-labelled CTP (Agilent Technologies), and hybridized onto SurePrint G3 Rat GE 8 x 60K Microarrays v2 (AMADID 074036; Agilent Technologies) for 16-20 h at 65°C in an Agilent oven with rotisserie. Following hybridization, the arrays were washed and scanned to derive the array images, using an Agilent microarray scanner G2505C. Feature Extraction Software (v11.5; Agilent) was used to generate the array data from the images, using Agilent grid 074036_D_F_20181024.

### Microarray Data Analysis

Data analysis was performed in R^48^ using the LIMMA package^49^. Grid annotation for the AMADID 074036 microarray was downloaded from Agilent eArray, and gene annotations were obtained from the Rattus.norvegicus R package based on the M5 genome build^50^. Microarray data were background corrected using the “normexp” method (with an offset of 50) and quantile normalized, producing normalized expression values in the log base 2 scale. Data were then filtered to remove all control probes, and non-control probes exhibiting low signals across the array sets. Keeping only those non-control probes that were at least 10% brighter than negative control probe signals on at least four arrays (based on the existence of four replicates per condition) left 31,691 out of 62,976 probes. Data from identical replicate probes within the same array were then averaged to produce expression values at the unique probe level, obtaining 20,326 unique probes, of which 15,247 are assigned to genome features with an Entrez gene identifier (Entrez ID).

Testing for differential expression between the experimental conditions was performed in LIMMA using linear modelling. Significance p-values were corrected for multiplicity using the Benjamini and Hochberg (BH) method, obtaining adjusted p-values (adj.P.Val). All the 20,326 unique probes were subjected to differential expression analysis but only the data for the significantly differentially expressed probes mapping to an Entrez ID (5,565) are reported in Supplementary File S1 and were considered in the downstream analyses.

Hierarchical clustering was performed by complete linkage clustering and using the Pearson correlation for the distance metric. Abundance values for each probe were transformed to z-scores for clustering. Functional enrichment analysis was performed using the R package clusterProfiler (version 3.8.1)^51^. The software produces adjusted p-values (p.adjust) using the BH correction method.

### Data Availability

Raw data in standard format from the microarray experiments have been deposited in ArrayExpress (www.ebi.ac.uk/arrayexpress/) with Accession number E-MTAB-8098.

## Acknowledgments

We gratefully acknowledge the financial support from Prof Matteo Santin of the Centre for Regenerative Medicine and Devices, University of Brighton.

## Author Contributions

CPS and PG conceived the study and supervised the project, MM and GB carried out the experiments, AH analyzed the results, MM, AH and PG wrote the manuscript although all the authors contributed to data analysis and to elaboration of the figures and of the final manuscript.

## Additional Information

The authors declare that they have no competing interest.

