## Supplementary Figure S1 and Supplementary Figure S2 for "Vitamins D3 and D2 have marked but different global effects on gene expression in a rat oligodendrocyte precursor cell line"

& Colin P. Smith

This PDF file includes:

Supplementary Figure 1**.** Functional enrichment analysis of the genes down-regulated by vitamin D.

Supplementary Figure 2. Functional enrichment analysis of the genes up-regulated by vitamin D.

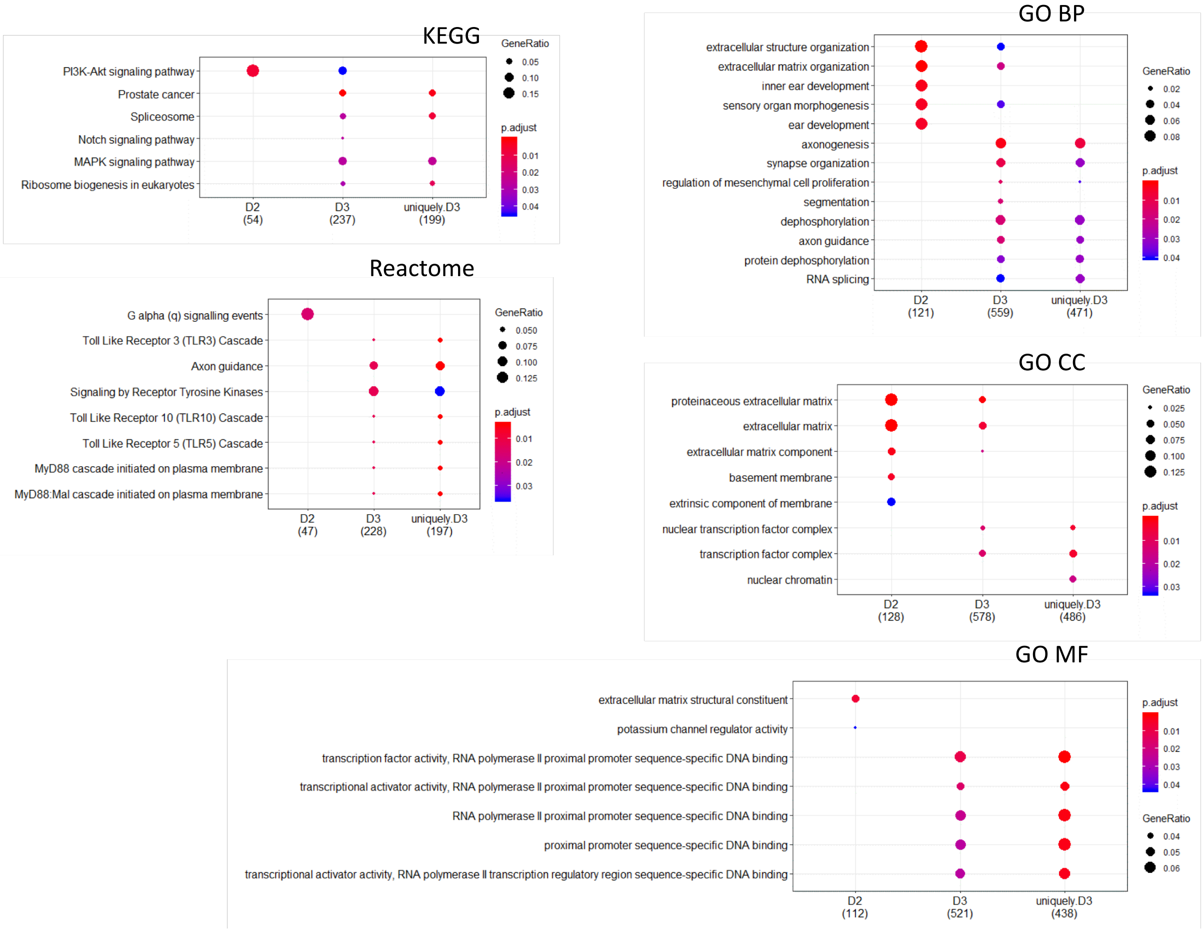


**Supplementary Figure S1**. **Functional enrichment analysis of the genes down-regulated by vitamin D.** Comparative functional enrichment analysis of the genes corresponding to the probes significantly decreasing in abundance after 24 h in response to either vitamin D2 (D2) or D3 (D3) and of those uniquely downregulated by D3 (uniquely.D3). Only the top significant categories are shown. Full results are provided in Supplementary File S3.

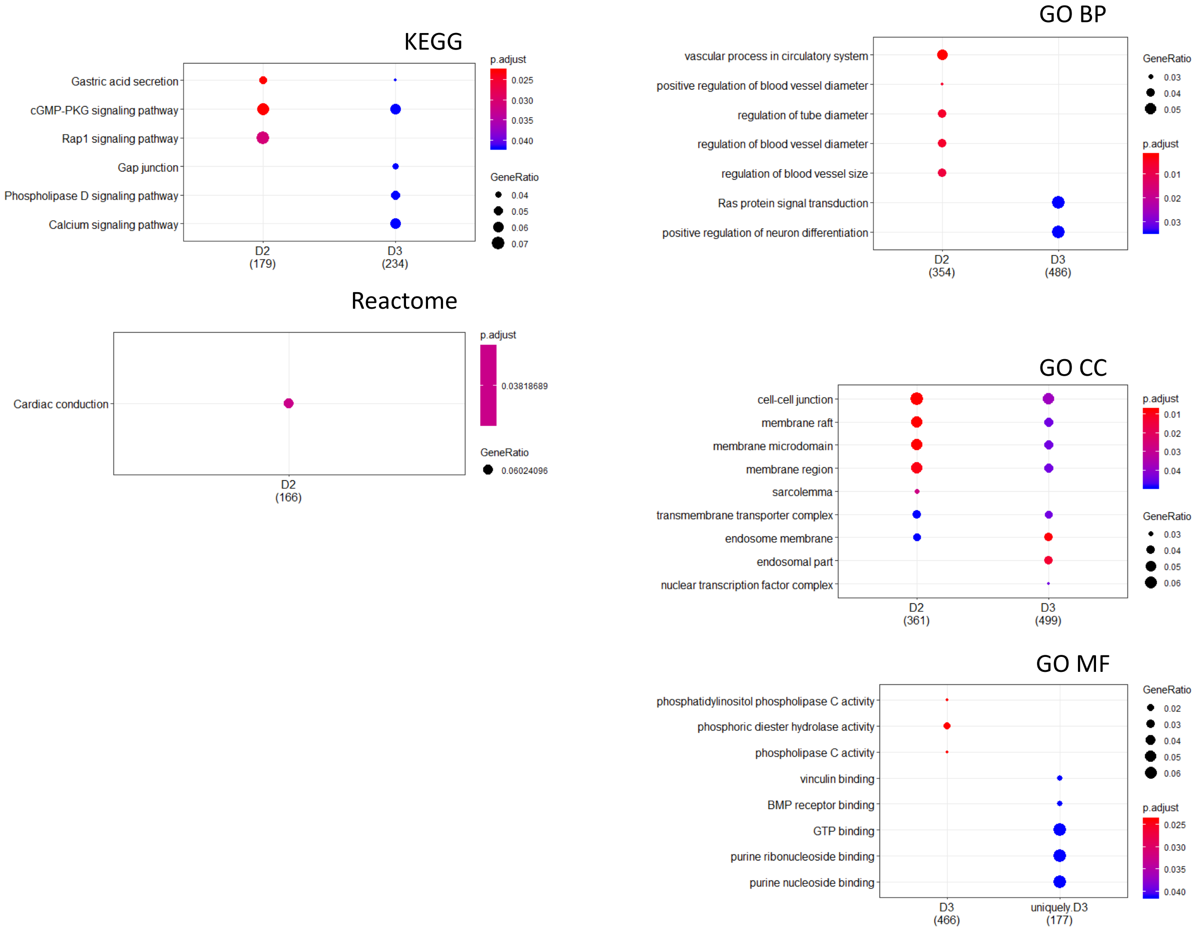


**Supplementary Figure S2**. **Functional enrichment analysis of the genes up-regulated by vitamin D.** Comparative functional enrichment analysis of the genes corresponding to the probes significantly increasing in abundance after 24 h in response to either vitamin D2 (D2) or D3 (D3) and of those uniquely downregulated by D3 (uniquely.D3). Only the top significant categories are shown. Full results are provided in Supplementary File S3.
